# Individual variability in polyphenol metabolite production: A dietary challenge study

**DOI:** 10.1101/2025.03.20.644335

**Authors:** Jessica K Sprinkles, Anju Lulla, Katie A Meyer

**Author notes:** **Corresponding Author:** Katie A Meyer, ScD, MPH, UNC Nutrition Research Institute, 500 Laureate Way, Suite 3201, Kannapolis, NC 28081.

## Abstract

**Background:** Polyphenols are metabolites of a variety of foods, mostly plant-based, that possess beneficial effects on metabolic health, largely through antioxidant and anti-inflammatory mechanisms. Beneficial associations between polyphenol-rich foods and metabolic health are well established, but individual differences in polyphenol metabolism exist and contribute to inconsistent relationships between plasma polyphenols and health outcomes. Characterizing differences in urinary polyphenol metabolite production may hold implications for studies of urinary polyphenol concentrations and advance understanding of the heterogeneity that exists in response to dietary interventions.

**Methods:** Eligible participants were recruited from the Kannapolis, NC area and underwent a controlled dietary challenge. Participants collected time-stamped urine samples for 24-hours after consumption of capsules containing soy isoflavone extract, flaxseed hull extract, pomegranate juice, and choline bitartrate. Urine samples were kept in coolers for delivery to the Nutrition Research Institute (NRI). Prior to the challenge, participants consumed meals low in polyphenols designed by the Metabolite core. Mass spectrometry quantified polyphenol metabolites from time-stamped urine samples. Metabolite concentrations were transformed to area under the curve (AUC) and maximum output variables, for total urine and 8-hr increments. Participants were grouped by tertile of metabolite output for each variable. Agreeance between metabolite output tertiles for each variable was assessed to identify consistency in measures.

**Conclusions:** In a likely homogenous sample of adults, we noted significant variability in polyphenol metabolite production in response to a controlled dietary challenge. In addition, there was generally low agreement between tertiles of metabolite output variables, holding implications for studies using time-specific metabolite measures, rather than 24-hr AUC.

## INTRODUCTION

Polyphenols comprise a diverse class of aromatic compounds, including flavonoids, phenolic acids, lignans, and stilbenes^1^. Their high concentration in plants and plant-based foods motivate the hypothesis that they contribute to the robust finding that chronic disease risk is inversely associated with dietary intake of fruits, vegetables, grains, and nuts and seeds^2-11^. However, despite their hypothesized importance, findings from human studies have been inconsistent because it is challenging to rigorously study the direct health effects of polyphenols due to their complex metabolism.

One factor contributing to the complex metabolism of polyphenols is the wide range of metabolites produced. More than 500 polyphenols have been identified with differing bioactivity and bioavailability due to culinary methods, habitual diet, or consumption of other compounds that may interfere with phase I or II metabolism^12,13^. Metabolism of polyphenols is dependent on several enzymatic reactions, gut microbiota, absorption and transport, all of which can affect bioactivity of these compounds^13-16^. As a result of this, high variability of polyphenol metabolite production and excretion has been documented in the literature^13,15^. To understand this variability, phenotypes of metabolite production, or metabotypes, of polyphenol metabolism have been suggested^17^. As such, production and excretion of polyphenol metabolites might not directly reflect polyphenol intake, making associations between polyphenols and disease challenging to study. While many studies are unable to correlate the beneficial effects of polyphenol intakes with circulating polyphenols, few studies have focused on patterns of polyphenol metabolite production and excretion following a controlled diet challenge^14^. This information is needed to determine the prevalence of metabolite production, define time periods for sample collection, and identify sources of variability in metabolite production.

In this study, we addressed the specific question of whether we could identify a time frame for collection of a single spot urine after controlled intake of polyphenol dietary sources to increase validity and decrease participant burden in epidemiological studies of polyphenol-disease associations. Currently, in large-scale epidemiological studies, a single blood sample and spot urine are collected from participants in the fasted state. The extent to which polyphenol metabolites will be detected in these spot urine or single blood samples is subject to the individual’s previous dietary intake and metabolic differences. We propose that methods to distinguish individuals with respect to metabolite production would enhance our ability to study the dietary effects of polyphenol sources, akin to a gene-nutrient interaction study. Here we used a combined dietary challenge in 40 participants to assess polyphenol metabolite excretion over a 24-hr urinary collection period.

## METHODS

### Study sample

We recruited 40 individuals from the Kannapolis, NC region. Recruitment was through advertisements on campus and in community venues (e.g., YMCA), as well as through outreach via a listserv maintained by the Nutrition Research Institute (NRI). Participants contacted the study recruiter through email to schedule a phone call to assess eligibility and confirm interest in taking part. Eligibility criteria included: 1) aged 18-70, 2) 18 ≤ BMI ≤ 32, 3) no weight gain or loss greater than 10% of their current body weight in the past year, 4) not pregnant or breastfeeding, 5) no known food allergies or dietary restrictions, 6) comfort swallowing pills, 7) ≤ 2 alcohol beverages per day, 8) non-smoker, 9) no antibiotics in the past 3 months, 10) no inflammatory bowel disease or other major chronic disease, 11) not taking chemotherapeutics or immune-suppressants, and 12) no history of major abdominal surgery (bowel resection, bariatric surgery).

### Challenge design

In a single-arm human intervention study, all 40 subjects consumed, in a single administration, on-site at the NRI: 2 commercial capsules of cocoa extract, 2 commercial capsules of soy isoflavone extract, 2 commercial capsules of flaxseed hulls extract, 8 ounces POM Wonderful pomegranate juice, and 1 commercial capsule of choline bitartrate.

Subjects were instructed to collect all urine for the 24 hours following the challenge, with each void in separate collection containers. Urine was collected in separate h-urine at their needs, so time collection points were different among subjects. Participants were provided coolers with gel packs to store samples until delivery to the NRI. Participants were provided meals for 2 days before the challenge and the day of the challenge to standardize diet across participants and to limit intake of metabolite precursors prior to the challenge. Meals were designed and prepared by a research dietitian.

### Urine sample preparation

Urine samples were defrosted, vortexed, and an aliquot of 250 µL of urine were added with 750 µL of 0.1% formic acid in water. Urine samples were vortexed for 1 min, centrifuged at 13,765 x g for 10 min at 4 °C, filtered and supernatants were finally analyzed by UHPLC-ESI-MS/MS.

### UHPLC/MS Analysis

Biological extracts were analyzed using an UHPLC DIONEX Ultimate 3000 equipped with a triple quadrupole TSQ Vantage (Thermo Fisher Scientific Inc., San José, CA, USA) fitted with a heated-ESI (H-ESI) (Thermo Fisher Scientific Inc., San José, CA, USA) probe. Separations were carried out by means of a Kinetex EVO C18 (100 × 2.1 mm), 2.6 μm particle size (Phenomenex, Torrance, CA, USA). Volume injection was 5 µL and column oven was set to 40°C. Compound elution was performed at a flow rate of 0.4 mL/min. For separation, mobile phase A was 0.2% formic acid in water and mobile phase B was acetonitrile containing 0.2% formic acid. The gradient started with 5% B, keeping isocratic conditions for 0.5 min, reaching 95% B at 7 min, followed by 1 min at 95% B and then 4 min at the start conditions to re-equilibrate the column. The applied mass spectrometry (MS) method consisted in the selective determination of each target precursor ion by the acquisition of characteristic product ions in selective reaction monitoring (SRM) mode, applying negative ionization mode. The spray voltage was set at 3 kV, the vaporizer temperature at 300 °C, and the capillary temperature operated at 270 °C. The sheath gas flow was 50 units, and auxiliary gas pressure was set to 10 units. Ultrahigh purity argon gas was used for collision-induced dissociation (CID). The S-lens values were defined for each compound based on infusion parameter optimization (Table 1). Conversely, for compounds that were not available for infusion, the S-lens values were set using the values obtained for the chemically closest available standards. Quantification was performed with calibration curves of standards, when available, or using the most structurally similar compound. Data processing was performed using Xcalibur software (Thermo Scientific Inc., Waltham, MA, USA).

**Table 1.**
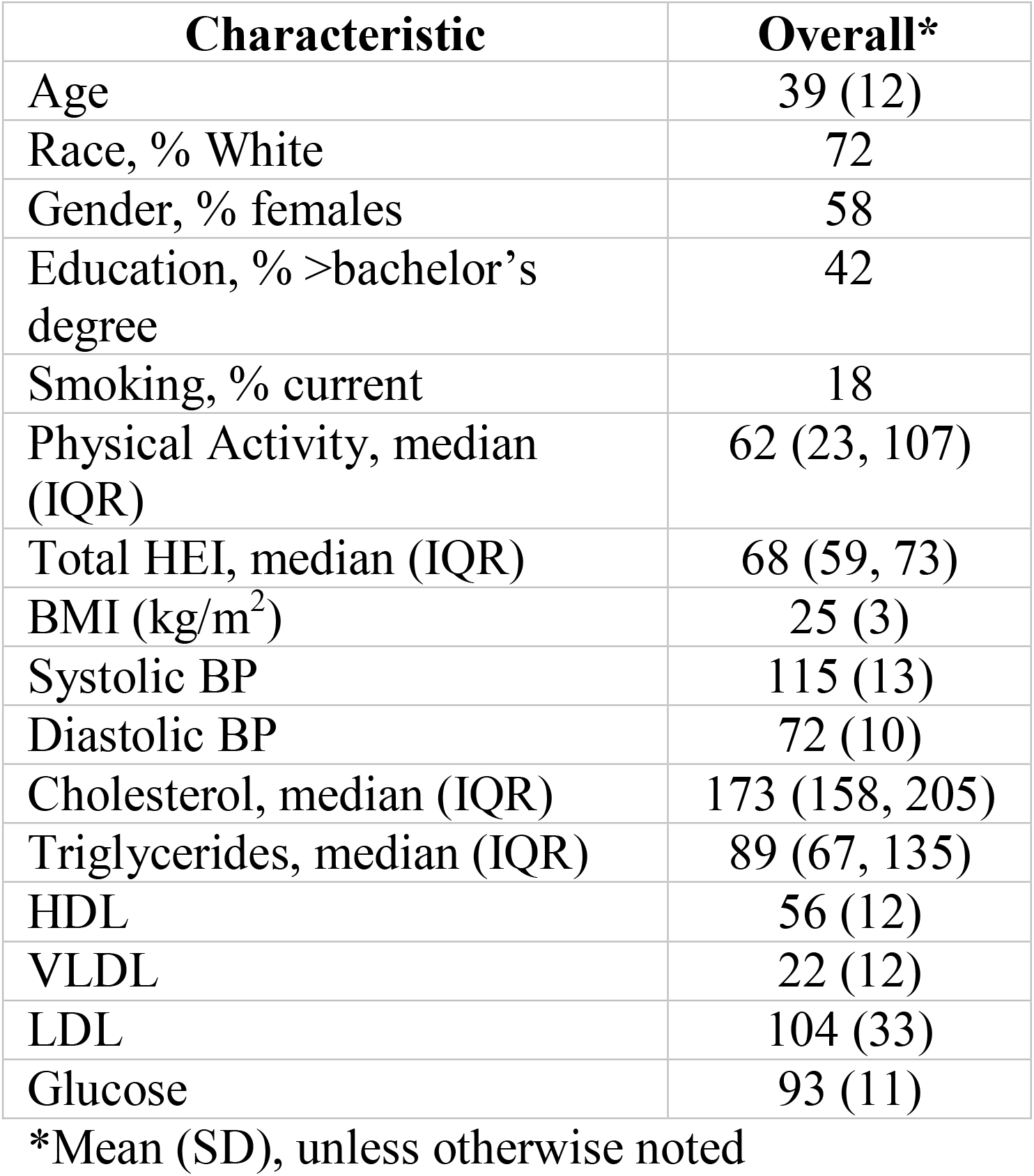
Participant characteristics of the study sample (n=40)

| **Characteristic** | **Overall\*** |
| --- | --- |
| Age | 39 (12) |
| Race, % White | 72 |
| Gender, % females | 58 |
| Education, % >bachelor's degree | 42 |
| Smoking, % current | 18 |
| Physical Activity, median (IQR) | 62 (23, 107) |
| Total HEI, median (IQR) | 68 (59, 73) |
| BMI (kg/m<sup>2</sup>) | 25 (3) |
| Systolic BP | 115 (13) |
| Diastolic BP | 72 (10) |
| Cholesterol, median (IQR) | 173 (158, 205) |
| Triglycerides, median (IQR) | 89 (67, 135) |
| HDL | 56 (12) |
| VLDL | 22 (12) |
| LDL | 104 (33) |
| Glucose | 93 (11) |

\*Mean (SD), unless otherwise noted

### Statistical Analysis

We transformed metabolite concentrations into six different variables: raw area under the curve (AUC), creatinine normalized, volume normalized, maximum, 8-hr increment AUC, and 8-hr increment maximum concentrations. We referred to the total AUC as the standard method for analyzing metabolite concentration data. Creatinine variables were AUCs normalized by mM of creatinine to produce uM of metabolite/mM of creatinine. Volume variables were AUCs (uM of metabolite) divided by the volume (L) of urine in samples. Maximum variables were the maximum output reached during the 24-hr collection period. Lastly, we calculated 8-hr increment (0-8hr, 8-16hr, 16-24hr) AUC and maximum values.

We generated descriptive characteristics for the overall study population. We displayed participants metabolite production over time in line graphs to assess individual output variability. Metabolite area under the curves (AUC) were summarized with medians (IQR) and % of non-producers. Phenotypes of non-producing ellagitannin and lignan sulfate metabolites were assessed using Venn diagrams of each subclass. We summarized participants’ time to reach maximum output of each metabolite with medians (IQR). All metabolite variables were grouped into tertiles. Agreeance in participant tertile classification between metabolite variables was assessed using alluvial plots and % misclassification. All statistical analysis was conducted in R (version 4.2.3) and metabolite output figures were created in Tableau.

## RESULTS

### Participant characteristics and variability

Study participants were generally healthy, as recruited, with average BMI’s, blood pressure, lipids, and glucose within normal ranges (Table 1). Timing and AUC of metabolite production was highly variable in the study sample (Table 2 and Figure 1). Time to reach maximum urinary output of metabolites varied across participants and metabolites (Table 3).

**Table 2.** Total (AUC) polyphenolic metabolite productions.

| Metabolite | AUC Median (IQR) | % Non-producers |
| --- | --- | --- |
| *Ellagitannin metabolites (phenolic acids) – pomegranates* |  |  |
| Urolithin A 8'-glucuronide | 4,102 (84, 20,496) | 22 |
| Isourolithin A 3'-glucuronide | 6,492 (1,800, 16,810) | 5 |
| Urolithin B 3'-glucuronide | 6,109 (3,840, 10,219) | 7.5 |
| *Isoflavone metabolites (flavonoids) - Soy* |  |  |
| Equol 7'-glucuronide | 2,158 (988, 8,389) | 0 |
| *Lignans - flaxseed* |  |  |
| Enterodiol glucuronide | 34,208 (17,828, 55,779) | 0 |
| Enterolactone glucuronide | 85,743 (35,914, 214,243) | 0 |
| Enterodiol sulfate | 2,248 (0, 5,455) | 30 |
| Enterolactone sulfate | 532 (0, 2,265) | 30 |
| *Valerolactones* |  |  |
| 5-Phenyl-y-valerolactone-3'-sulfate | 246,307 (112,432, 522,357) | 0 |
| 5-(Hydroxyphenyl)-g-valerolactone-sulfate | 7,756,073 (2,493,575, 18,169,349) | 0 |
| 5-(5'-Hydroxyphenyl)-y-valerolactone-3'-sulfate | 16,391 (4,074, 46,381) | 0 |
| 5-Phenyl-y-valerolactone-3'-glucuronide | 16,682 (9,796, 52,624) | 2.5 |
| 5-(3'-Hydroxyphenyl)-y-valerolactone-4'-glucuronides | 166,336 (44,734, 670,814) | 0 |
| 5-(4'-Hydroxyphenyl)-y-valerolactone-3'-glucuronide | 39,590 (8,303, 140,057) | 0 |
| 5-(5'-Hydroxyphenyl)-y-valerolactone-3'-glucuronide | 2,413 (1,311, 8,250) | 0 |

**Table 3.**
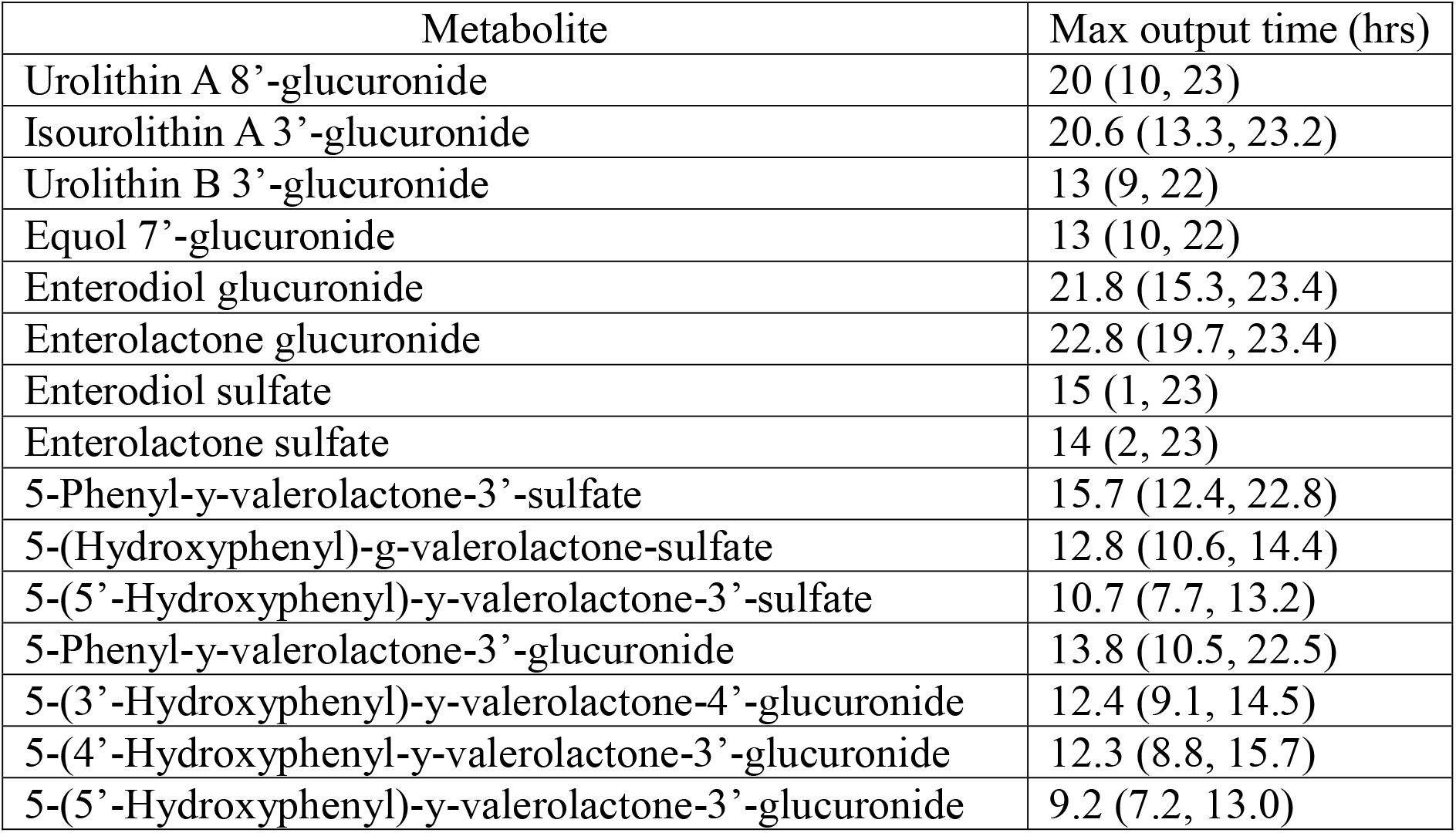
Summary statistics of time that max metabolite output was reached.

**Figure 1.**
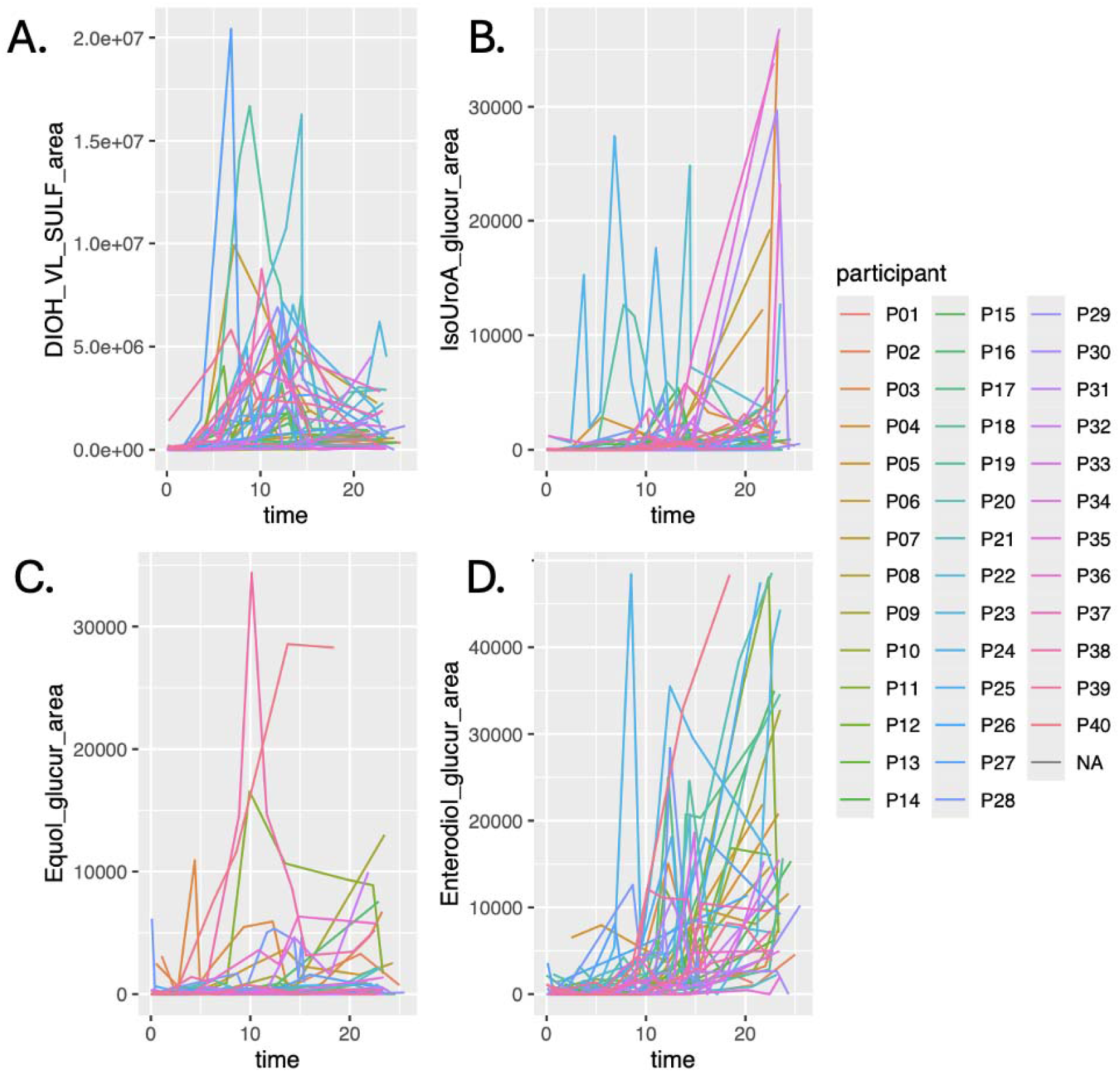
Individual variability in metabolite production over time for A. 5-(Hydroxyphenyl)-g-valerolactone-sulfate, B. Isourolithin A 3’-glucuronide, C. Equol 7’-glucuronide, and D. Enterodiol Glucuronide.

### Observed non-producer phenotypes

We observed non-producers of many urinary metabolites. Of the study sample, 30% were non-producers of enterodiol sulfate and enterolactone sulfate, 22% were non-producers of urolithin A 8’-glucuronide, 7.5% were non-producers of urolithin B 3’-glucuronide, 5% were non-producers of isourolithin A 3’-glucuronide, and 2.5% were non-producers of 5-phenyl-y-valerolactone-3’-glucuronide (Table 2). Urolithin A 8’-glucuronide, enterodiol sulfate, and enterolactone sulfate were most commonly not produced together (n=4), followed by non-producers of both enterodiol sulfate and enterolactone sulfate (n=3), and non-producers of both enterolactone sulfate and urolithin A 8’-glucuronide (n=2) (Figure 2). Majority of participants (70%) produced all three ellagitannin metabolites (urolithin A 8’-glucuronide, isourolithin A 3’-glucuronide, and urolithin B 3’-glucuronide). 17.5% of participants did not produce urolithin A 8’-glucuronide but were producers of the other two ellagitannin metabolites. 7.5% of participants did not produce urolithin B 3’-glucuronide but were producers of the other two ellagitannin metabolites. 5% of participants produced only urolithin B 3’-glucuronide out of the ellagitannin metabolites (Figure 3A). For lignans, all participants produced enterodiol glucuronide and enterolactone glucuronide, but 30% of participants did not produce enterodiol sulfate and/or enterolactone sulfate. 57.5% of participants were producers of both enterodiol- and enterolactone-sulfate, while 12.5% participants produced only enterodiol sulfate and 12.5% of participants produced only enterolactone sulfate (Figure 3B).

**Figure 2.**
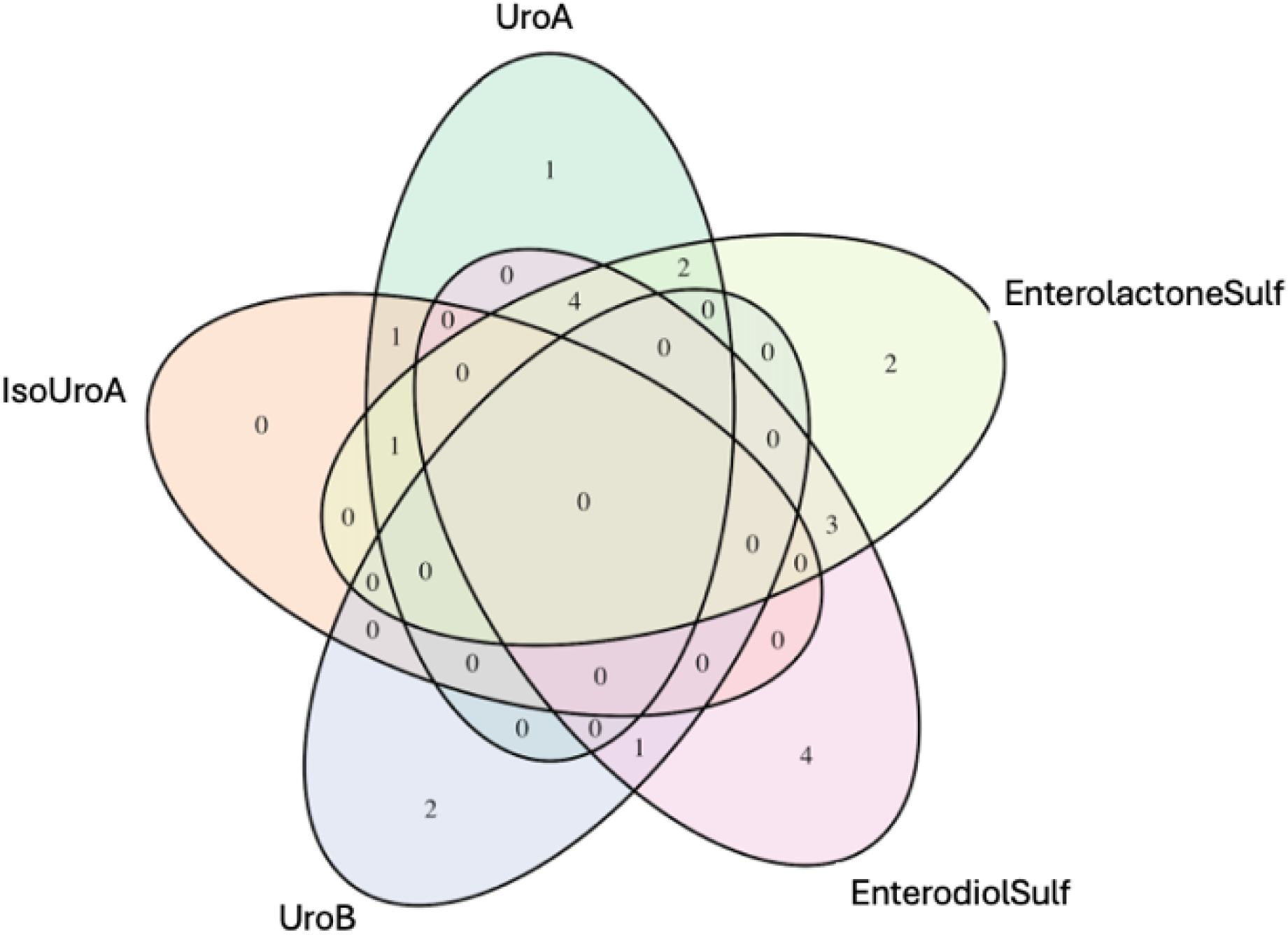
Venn diagram of non-produced ellagitannin (Urolithin A Glucuronide, IsoUrolithin A Glucuronide, and Urolithin B Glucuronide) and lignan (Enterolactone sulfate and Enterodiol sulfate) metabolites.

**Figure 3.**
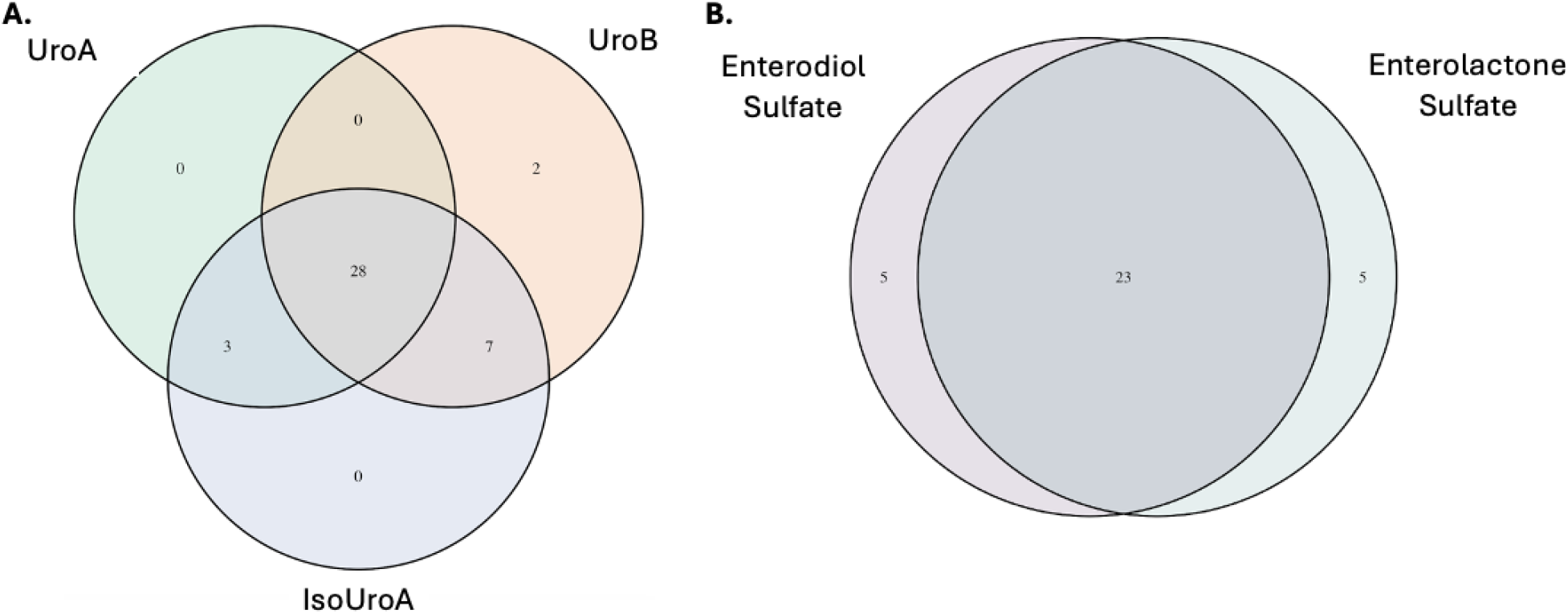
Venn diagram of A. ellagitannin metabolite producers and B. lignan producers.

### Participant classification by metabolite variables

Next, we assessed differences in the ranking of participants into producer tertiles (low, medium, and high) by the measure of metabolite production used (ex. AUC vs. 24-hr maximum). Classification of participants as low, medium, and high metabolite producers did differ depending on the measure used. Across all metabolites and measures used, % of participants that were classified differently from the AUC classification varied from 5%-68%. AUCs and maximums of each 8-hour increment during the 24-hr collection period showed high percentages of misclassification compared to total AUC tertiles (Table 4). In alluvial plots comparing total AUC, creatinine-adjusted, 24-hr maximum, and volume-adjusted tertiles, the extent of changing tertile classifications also varied by metabolite. For example, urolithin B 3’-glucuronide had significant changes in participant classification comparing the creatinine adjusted tertiles to the maximum output tertiles (Figure 4A). Other metabolites, like 5-Phenyl-y-valerolactone-3’-sulfate showed consistent tertile classifications between the AUC and creatinine adjusted variable (Figure 4D). Maximum variables appeared to contribute to the most misclassification across many metabolites.

**Table 4.** Misclassification (%) of participants by concentration variable used (high vs. mid vs. low producer: tertiles) compared to total AUC classification.

|  | Volume<br>Variable<br>(n=40) | Creatinine<br>Variable<br>(n=40) | Overall<br>Max<br>(n=40) | 0-8hr<br>Max<br>(n=40) | 0-8hr<br>AUC<br>(n=40) | 8-16hr<br>Max<br>(n=40) | 8-16hr<br>AUC<br>(n=40) | 16-24hr<br>Max<br>(n=37) | 16-24hr<br>AUC<br>(n=37) |
| --- | --- | --- | --- | --- | --- | --- | --- | --- | --- |
| *Ellagitannin metabolites - pomegranates* |  |  |  |  |  |  |  |  |  |
| Urolithin A glucuronide | 5 | 5 | 20 | 48 | 48 | 25 | 25 | 24 | 19 |
| Isourolithin A glucuronide | 20 | 20 | 10 | 48 | 48 | 45 | 40 | 19 | 24 |
| Urolithin B glucuronide | 40 | 25 | 30 | 50 | 50 | 32 | 30 | 32 | 43 |
| *Isoflavone metabolites – Soy* |  |  |  |  |  |  |  |  |  |
| Equol 7'-glucuronide | 25 | 20 | 10 | 50 | 42 | 40 | 42 | 14 | 14 |
| *Lignans – flaxseed* |  |  |  |  |  |  |  |  |  |
| Enterodiol-glucuronide | 48 | 35 | 35 | 70 | 70 | 45 | 45 | 43 | 32 |
| Enterolactone-glucuronide | 30 | 30 | 30 | 50 | 40 | 25 | 30 | 32 | 22 |
| Enterodiol-sulfate | 20 | 10 | 25 | 55 | 55 | 40 | 40 | 22 | 22 |
| Enterolactone-sulfate | 10 | 5 | 5 | 42 | 42 | 28 | 28 | 22 | 22 |
| *Valerolactones* |  |  |  |  |  |  |  |  |  |
| 5-Phenyl-y-valerolactone-3'-sulfate | 20 | 10 | 25 | 62 | 60 | 22 | 18 | 43 | 41 |
| 5-(Hydroxyphenyl)-g-valerolactone-sulfate | 20 | 5 | 20 | 40 | 50 | 30 | 25 | 49 | 43 |
| 5-(5'-Hydroxyphenyl)-y-valerolactone-3'-sulfate | 15 | 10 | 10 | 15 | 25 | 20 | 10 | 30 | 41 |
| 5-Phenyl-y-valerolactone-3'-glucuronide | 15 | 15 | 20 | 65 | 68 | 20 | 20 | 38 | 41 |
| 5-(3'-Hydroxyphenyl)-y-valerolactone-4'-glucuronide | 30 | 15 | 10 | 38 | 38 | 10 | 10 | 38 | 46 |
| 5-(4'-Hydroxyphenyl)-y-valerolactone-3'-glucuronide | 15 | 15 | 10 | 30 | 30 | 10 | 15 | 41 | 41 |
| 5-(5'-Hydroxyphenyl)-y-valerolactone-3'-glucuronide | 30 | 40 | 15 | 15 | 20 | 30 | 20 | 43 | 43 |

**Figure 4.**
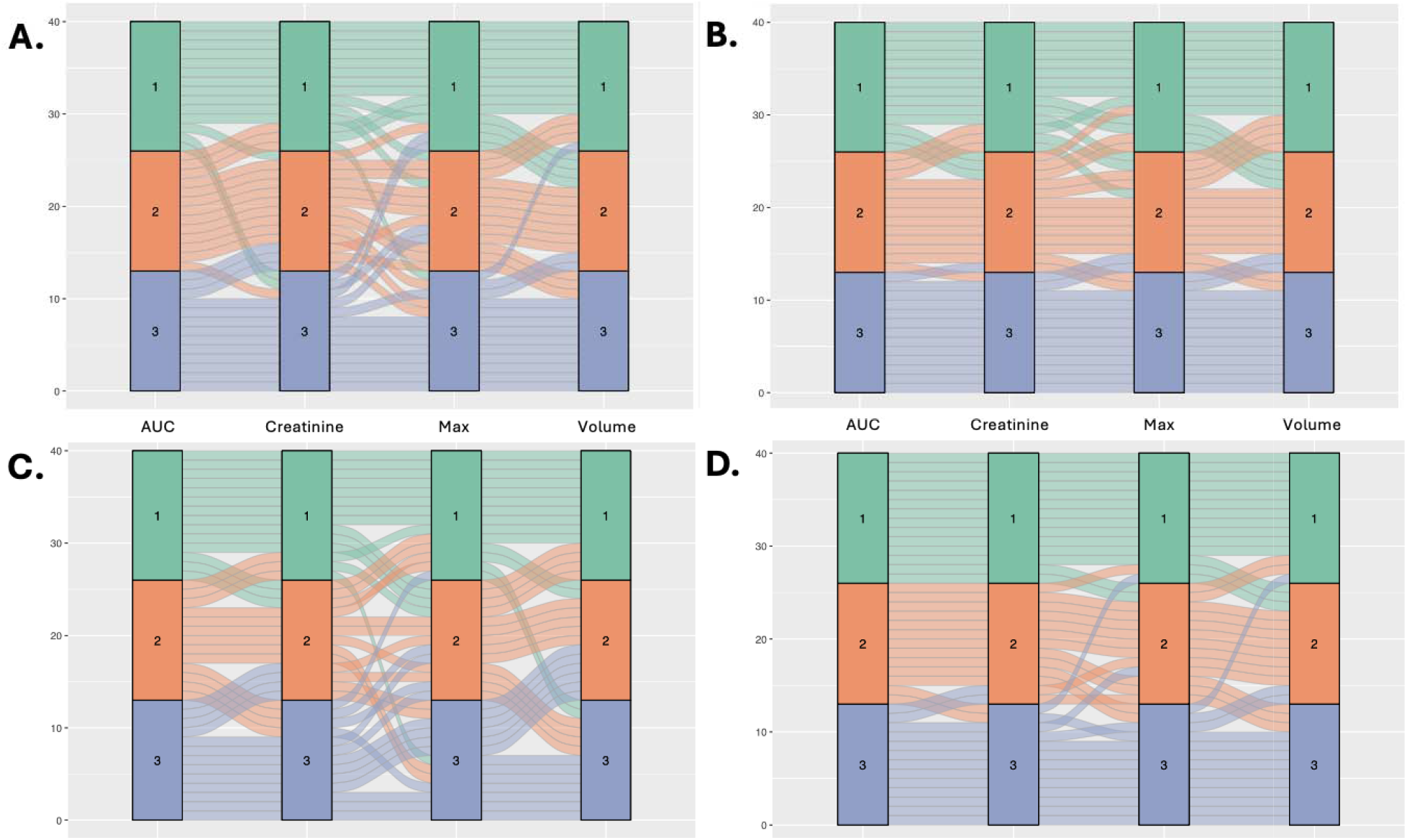
Alluvial plots of tertile classification agreement between AUC, max, volume, and creatinine variables for A. Urolithin B 3’-glucuronide, B. Equol Glucuronide, C. Enterodiol Glucuronide, and D. 5-Phenyl-y-valerolactone-3’-sulfate.

## DISCUSSION

In this dietary challenge study of generally healthy adults, we observed significant heterogeneity in polyphenol metabolite output in response to foods rich in polyphenols. Significant inter-individual variability in urinary polyphenol metabolites has been demonstrated in previous studies of free-living participants and been contributed to many variables including polyphenol bioavailability and bioactivity, sex, genetics, gut microbiota, ethnicity, age, and habitual dietary intake^18,19^. In addition, urinary metabolites may serve as biomarkers of intake for some polyphenol metabolites, but not others^21^. In our study, we observed significant variability in metabolite production in a largely homogenous and generally healthy sample of adults under a controlled diet challenge.

Variability seen in polyphenol metabolism has prompted recent literature to determine polyphenol metabotypes, or phenotypes of polyphenol metabolite production, but our results are inconsistent with previously identified metabotypes^17^. Metabotypes of ellagitannins have been observed in previous human intervention trials: phenotype A produces only urolithin A; phenotype B produces urolithin A and one, or both, of isourolithin A and urolithin B; phenotype O does not produce any of these three metabolites^20^. We primarily observed phenotype B (n=28) rather than phenotype A, which has been the predominant metabotype observed in other studies, and did not observe phenotype A or O (Figure 3A)^20^. Reasons for this are unclear, as these previous intervention studies were also collecting 24hr urine samples in generally healthy male and female participants.

Urinary excretion of metabolites may be subject to differences in gut microbiota, but our data did not identify significant associations between gut microbiota and metabolites^21^. Other variables may be contributing to variability in polyphenol metabolite production including variability that have been identified as confounders of polyphenol bioavailability and absorption including age, habitual energy intake, sex, and BMI^15^. Larger controlled diet studies of polyphenol metabolism in diverse populations are needed to detect associations between participant characteristics and metabolite output.

This current study identified significant variability in metabolite production in response to a controlled diet challenge. A strength of this study is that participants collected timed urinary samples, allowing for time-specific analysis of excretion, in addition to a pooled 24-hr urinary concentration. We additionally were able to control participants’ recent dietary intake to minimize the influence of recent habitual diet.

Previous studies have suggested that 24-hr urine measures are better indicators of polyphenol intake than spot measures, but 24-hr urinary collection is burdensome^15^. Ideally, a single spot urine sample would be able to accurately rank participants total metabolite production. Our data revealed that there is high variability in the time it takes for participants to reach maximum metabolite output and that some participants have greater maximum peaks than others, regardless of 24-hr AUC. In analysis of classification of metabolite producer tertiles, we saw high percentages of tertile reclassification for 0-8hr, 8-16hr, and 16-24hr measures compared to the 24-hr AUC. Given these results, we propose that single spot urine samples are not able to accurately reflect polyphenol intake or metabolism and that epidemiological studies of single spot urine samples are subject to misclassification.

## References

1. Han X, Shen T, Lou H. Dietary Polyphenols and Their Biological Significance. International Journal of Molecular Sciences. 2007;8(9):950–988.

2. Panche AN, Diwan AD, Chandra SR. Flavonoids: an overview. Journal of Nutritional Science. 2016;5:e47.

3. Sansone R, Rodriguez-Mateos A, Heuel J, et al. Cocoa flavanol intake improves endothelial function and Framingham Risk Score in healthy men and women: a randomised, controlled, double-masked trial: the Flaviola Health Study. British Journal of Nutrition. 2015;114(8):1246–1255.

4. Adriouch S, Lampuré A, Nechba A, et al. Prospective Association between Total and Specific Dietary Polyphenol Intakes and Cardiovascular Disease Risk in the Nutrinet-Santé French Cohort. Nutrients. 2018;10(11):1587.

5. Zamora-Ros R, Forouhi NG, Sharp SJ, et al. Dietary Intakes of Individual Flavanols and Flavonols Are Inversely Associated with Incident Type 2 Diabetes in European Populations1, 2, 3. The Journal of Nutrition. 2014;144(3):335–343.

6. Zamora-Ros R, Forouhi NG, Sharp SJ, et al. The Association Between Dietary Flavonoid and Lignan Intakes and Incident Type 2 Diabetes in European Populations: The EPIC-InterAct study. Diabetes Care. 2013;36(12):3961–3970.

7. Ding M, Pan A, Manson JE, et al. Consumption of soy foods and isoflavones and risk of type 2 diabetes: a pooled analysis of three US cohorts. European Journal of Clinical Nutrition. 2016;70(12):1381–1387.

8. Yang B, Chen Y, Xu T, et al. Systematic review and meta-analysis of soy products consumption in patients with type 2 diabetes mellitus. Asia Pacific Journal of Clinical Nutrition. 2011;20(4):593–602.

9. Pan A, Sun J, Chen Y, et al. Effects of a Flaxseed-Derived Lignan Supplement in Type 2 Diabetic Patients: A Randomized, Double-Blind, Cross-Over Trial. PLOS ONE. 2007;2(11):e1148.

10. Rhee Y, Brunt A. Flaxseed supplementation improved insulin resistance in obese glucose intolerant people: a randomized crossover design. Nutrition Journal. 2011;10:1–7.

11. Bakkalbaşi E, Menteş Ö, Artik N. Food Ellagitannins–Occurrence, Effects of Processing and Storage. Critical Reviews in Food Science and Nutrition. 2008;49(3):283–298.

12. Amiot MJ, Riva C, Vinet A. Effects of dietary polyphenols on metabolic syndrome features in humans: a systematic review. Obesity Reviews. 2016;17(7):573–586.

13. Castro-Barquero S, Lamuela-Raventós RM, Doménech M, Estruch R. Relationship between Mediterranean Dietary Polyphenol Intake and Obesity. Nutrients. 2018;10(10):1523.

14. Rajha HN, Paule A, Aragonès G, et al. Recent Advances in Research on Polyphenols: Effects on Microbiota, Metabolism, and Health. Molecular Nutrition & Food Research. 2022;66(1):2100670.

15. Clarke ED, Rollo ME, Collins CE, et al. The Relationship between Dietary Polyphenol Intakes and Urinary Polyphenol Concentrations in Adults Prescribed a High Vegetable and Fruit Diet. Nutrients. 2020;12(11):3431.

16. Serreli G, Deiana M. In vivo formed metabolites of polyphenols and their biological efficacy. Food & Function. 2019;10(11):6999–7021.

17. Tosi N, Favari C, Bresciani L, et al. Unravelling phenolic metabotypes in the frame of the COMBAT study, a randomized, controlled trial with cranberry supplementation. Food Research International. 2023;172:113187.

18. Mennen LI, Sapinho D, Ito H, Galan P, Hercberg S, Scalbert A. Urinary excretion of 13 dietary flavonoids and phenolic acids in free-living healthy subjects – variability and possible use as biomarkers of polyphenol intake. European Journal of Clinical Nutrition. 2008;62(4):519–525.

19. Cassidy A, Minihane A-M. The role of metabolism (and the microbiome) in defining the clinical efficacy of dietary flavonoids1. The American Journal of Clinical Nutrition. 2017;105(1):10–22.

20. Tomás-Barberán FA, García-Villalba R, González-Sarrías A, Selma MV, Espín JC. Ellagic Acid Metabolism by Human Gut Microbiota: Consistent Observation of Three Urolithin Phenotypes in Intervention Trials, Independent of Food Source, Age, and Health Status. Journal of Agricultural and Food Chemistry. 2014;62(28):6535–6538.

21. Pérez-Jiménez J, Hubert J, Hooper L, et al. Urinary metabolites as biomarkers of polyphenol intake in humans: a systematic review1234. The American Journal of Clinical Nutrition. 2010;92(4):801–809.

22. Scalbert A, Morand C, Manach C, Rémésy C. Absorption and metabolism of polyphenols in the gut and impact on health. Biomedicine & Pharmacotherapy. 2002;56(6):276–282.

